# Integrative Functional Genomics Identifies ARHGAP10 in the 4q31.2 Locus as a Novel Congenital Heart Disease and Ciliopathy Gene

**DOI:** 10.1101/2025.10.10.681676

**Authors:** Ewelina Sochaka, Svanna Hinson, Dave Shook, Doug DeSimone, Saurabh Kulkarni

## Abstract

Congenital heart disease (CHD) remains a major cause of pediatric morbidity and mortality, yet its genetic underpinnings are not fully understood. Two studies independently identified rare deletions in ARHGAP10 (GAP10), a Rho GTPase-activating protein located at 4q31.2 in individuals with heterotaxy and atrial septal defects (n=2 rare copy number variants (CNVs)), highlighting GAP10 as a new candidate CHD gene. While the 4q31.2 locus is implicated in CHD, the function of GAP10 has not been investigated. Here, using *Xenopus tropicalis* as a model, we demonstrate that *gap10* deletion disrupts morphogenetic movements critical for body axis extension, left-right organizer (LRO) formation, and ciliogenesis, leading to severe cardiac looping defects that closely mirror human CHD phenotypes. Moreover, *gap10* localizes to basal bodies of motile cilia in multiciliated cells, where it regulates basal body organization and apical actin enrichment by recruiting focal adhesion kinase (FAK) to specialized ciliary adhesion complexes. In summary, our findings implicate GAP10 as a clinically relevant, genetically supported, and functionally validated regulator of CHD and ciliogenesis, underscoring the power of integrative functional genomics in the discovery of rare disease genes.

## INTRODUCTION

Congenital heart disease (CHD) is the most common class of major birth defects, affecting roughly 1% of live births. While recent advancements in sequencing technologies have expanded our understanding of CHD pathogenesis, many patients still lack a definitive genetic diagnosis due to high genetic heterogeneity and incomplete annotation of disease genes, underscoring the need to identify novel genes and elucidate their roles in critical developmental processes^1–6^.

Genome-wide screens have shown that rare (<1 %) copy-number variants (CNVs) are important contributors to CHD^5–7^. Recurrent multi-gene deletions at 22q11.2 and 1q21.1, and single-gene losses such as the ∼0.5 Mb deletion removing GATA4 at 8p23.1, exemplify how both large and focal CNVs can cause CHD^8–12^. In this context, a novel deletion affecting *ARHGAP10* (also known as *GAP10*), a gene encoding a Rho GTPase-activating protein, was previously identified in a patient with heterotaxy syndrome (Figure 1A)^6^. Heterotaxy syndrome is a severe developmental disorder characterized by the abnormal arrangement of visceral organs along the left-right (LR) axis, leading to profound clinical consequences such as congenital heart defects and impaired organogenesis, and frequent disruption of mucociliary clearance caused by impaired motile cilia function^13–17^. A second patient with an atrial septal defect (ASD) also carried a rare CNV in *GAP10* (Figure 1A)^18^. Additionally, multiple individuals with large CNVs spanning *GAP10* frequently presented CHD phenotypes such as ASD, ventricular septal defects (VSD), and abnormal heart morphology, often associated with heterotaxy-related presentations (Table 1) (DECIPHER). However, the biological mechanisms linking GAP10 disruption to the observed clinical phenotype/s have remained unexplored.

**Figure 1.**
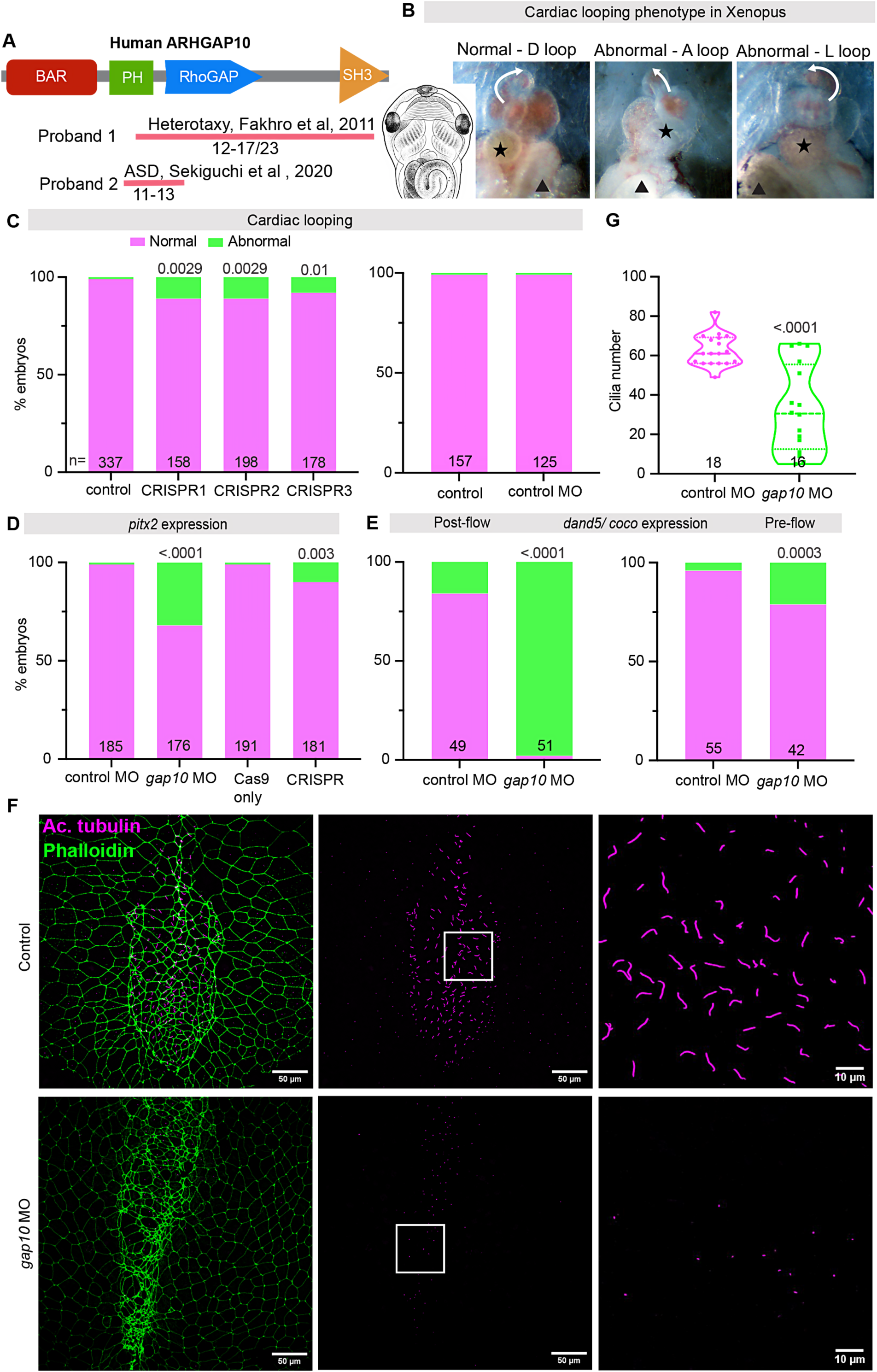
ARHGAP10 is Essential for Cardiac Looping and Left-Right Patterning in *Xenopus* Embryos. **(A)** Schematic of the domain structure of ARHGAP10, including BAR, PH, RhoGAP, and SH3 domains and locations of patient-specific CNVs. **(B)** Drawing (Zahn drawing) shows the position of internal organs in a wild-type embryo. Normal and abnormal cardiac looping phenotypes (white arrow) in *Xenopus* embryos. Black star and black arrowhead represent the position of the gallbladder and the gut. Note that the gallbladder is normally to the left of the embryo and the gut to the right side. However, in embryos with heart looping defects, their position can also be abnormal, indicating overall defects in left-right patterning. **(C)** % embryos with normal and abnormal cardiac looping phenotypes in wild-type, *gap10* CRISPRs, and control morpholino knockdown embryos. Quantification (bar graphs) shows proportion of normal (magenta) versus abnormal (green) heart looping (n indicated per group; p values shown). **(D)** Quantification of *pitx2* expression patterns (left-sided, abnormal – bilateral/right/absent) in controls and *gap10*-deficient embryos (n per group, p values as above). **(E)** Proportion of embryos with normal or abnormal coco/dand5 expression in the left-right organizer (LRO) at post-flow and pre-flow stages (n per group, p values as above). **(F)** Immunofluorescence images of LRO showing acetylated tubulin (magenta, cilia) and phalloidin (green, actin) in control and *gap10* morphant embryos, at low (scale bar: 50 μm) and high (10 μm) magnification. **(G)** Number of cilia in controls and *gap10*-deficient embryos (n per group, p values as above).

**Table 1.**
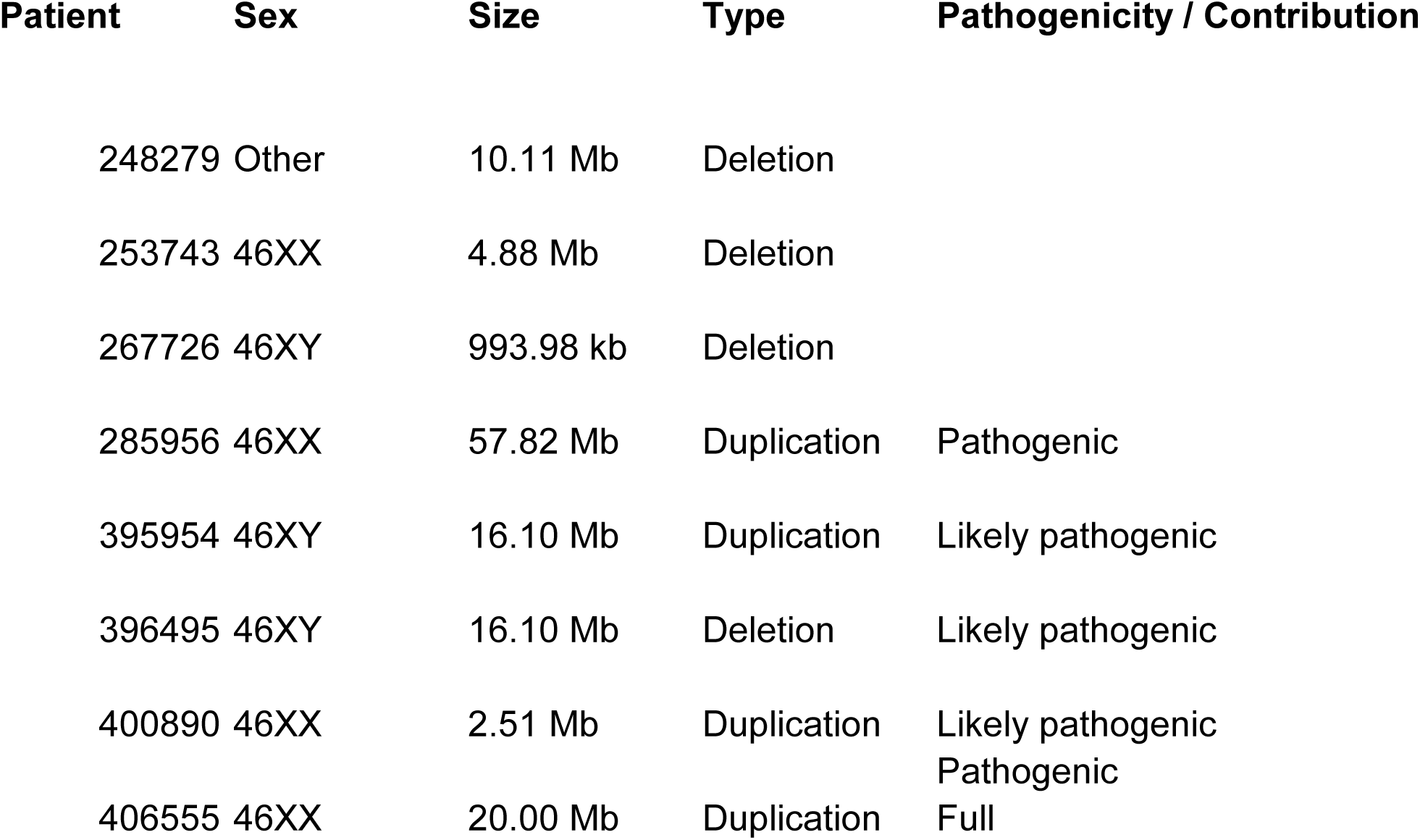
Clinical Phenotypes in Individuals with GAP10 (ARHGAP10) CNVs Identified in the DECIPHER Database. Summary of congenital heart disease and related phenotypes in patients with focal or large deletions spanning **ARHGAP10** at 4q31.2. Phenotypes include heterotaxy, atrial and ventricular septal defects, abnormal heart morphology, short stature, respiratory distress, and other developmental anomalies. CNV size, coordinates, inheritance, and DECIPHER IDs are indicated. See also Figure 1 for schematic CNV locations.

*GAP10* is a member of the RhoGAP superfamily involved in small GTPase signaling, a pathway implicated in CHD pathogenesis^18–21^. Notably, *GAP10* is located in the 4q31.2 locus near the EDNRA gene, a region implicated in CHD^22,23^. Importantly, *GAP10* is highly expressed in the embryonic heart and interacts with key regulators of the Rho signaling pathway.^18–21^ For example, focal adhesion kinase (FAK/PTK2), a direct interactor of GAP10, is essential for heart outflow tract formation and septation^19,21,24–27^. Pathogenic variants in other Rho regulators (e.g., ARHGAP31, DOCK6) yield syndromes that include structural heart malformations^28,29^. Together, these data suggest GAP10 as a plausible CHD candidate operating at the intersection of Rho-GTPase signaling, cytoskeletal remodeling, and LR patterning.

We hypothesize that ARHGAP10 is essential for cardiac development and that its disruption underlies the clinical spectrum observed in human patients. In this study, we employ comprehensive *in vivo* analyses, including CRISPR-mediated gene editing and morpholino (MO)-based knockdowns in *Xenopus* embryos, to provide compelling evidence that disrupting Gap10 causes significant cardiac looping defects, abnormal expression of LR patterning genes, and impaired embryonic growth. Additionally, our findings reveal that human GAP10 localizes specifically to the basal bodies of motile cilia, controlling the recruitment of focal adhesion kinase (FAK), basal body organization, and the actin cytoskeleton structure, all of which are essential for proper ciliogenesis and the function of motile cilia^30^. Overall, our research underscores the crucial role of GAP10 in pivotal developmental processes and, when considered alongside rare patient deletions, highlights its emerging importance as a genetically supported candidate gene for heterotaxy and CHD through an integrative functional genomics approach.

## RESULTS AND DISCUSSION

### *Gap10* is Essential for Left-Right Patterning and Cardiac Morphogenesis

We investigated the role of *gap10* in left-right (LR) axis formation using CRISPR-Cas9 gene editing in *Xenopus tropicalis*. Three independent single-guide RNAs (sgRNAs) targeting non-overlapping sites (two targeting exon 1 and one targeting exon 4) in *gap10* yielded consistent cardiac looping abnormalities (∼10%, Figure 1B, C). These abnormalities in *Xenopus* are a hallmark of disrupted LR patterning, as well as other associated CHD phenotypes^31–34^. Embryos with Gap10 knockdown via MO either died or showed severe developmental abnormalities, preventing analysis of heart looping. In contrast, control MO-injected embryos showed normal heart looping (Figure 1B, C). To assess LR molecular patterning, we examined *pitx2*, a canonical asymmetry marker normally expressed only in the left lateral plate mesoderm in *Xenopus* embryos^35^. Nearly all control MO or Cas9-only embryos exhibited the expected left-sided *pitx2* expression, whereas Gap10-deficient embryos frequently showed aberrant *pitx2* patterns, including bilateral, right-sided, or absent expression (Figure 1D). In normal development, leftward flow at the LR organizer (LRO) leads to suppression of the Nodal antagonist *dand5* (called *coco* in *Xenopus*) on the left side, thereby permitting left-sided *nodal* and *pitx2* activation^36–38^. We therefore examined *coco/dand5* at the LRO at stage 19 (post-flow). Control embryos exhibited the expected asymmetric *coco* expression (high on the right, suppressed on the left), while *gap10* morphants showed *coco* misexpression, such as persistent bilateral or left-sided *coco* expression (Figure 1E).

Because *coco/dand5* misexpression suggested a failure in cilia-driven LR signaling, we next analyzed LRO cilia and cell morphology in *gap10* morphants^32,38^. In controls, the LRO forms a characteristic teardrop-shaped pit covered with sensory and motile cilia (Figure 1F). *gap10* morphants exhibited an abnormally shaped LRO with clear cytoskeletal and ciliary defects: the LRO cilia were extremely short (appearing as punctate “dots” rather than the typical ∼5 μm hair-like projections) and markedly reduced in number (Figure 1F, G). Given the aberrant LRO morphology, we asked whether its initial patterning was disturbed. Prior to fluid flow (pre-flow, early gastrula, stage 16), *dand5/coco* is normally expressed symmetrically at the LRO margins^32,36^. In *gap10* morphants, *dand5* was already misexpressed before flow onset, indicating that LRO patterning was intrinsically disrupted (Figure 1E). However, the extent of *coco* misexpression dramatically increased from ∼30% of embryos before ciliary flow to nearly 90% after flow, indicating that a failure of nodal flow further exacerbates the LR patterning defects in *gap10* morphants. These data suggest that *gap10* is required for both proper LRO formation and motile cilia function in LR signaling. Consistent with the genetic link between GAP10 and heterotaxy, the spectrum of LRO defects in G*ap10*-deficient *Xenopus* embryos firmly establishes GAP10 as a critical gene in vertebrate LR patterning.

### *Gap10* Regulates Gastrulation and Body Axis Elongation During Embryogenesis

LRO formation is coupled to proper gastrulation, so we examined whether Gap10 loss impairs early morphogenetic movements^39^. Indeed, Gap10 depletion (by both morpholino and CRISPR) led to delayed or incomplete blastopore closure during gastrulation (Figure 2A-C). By tailbud stages, Gap10-deficient embryos developed severely shortened anteroposterior body axes (Figure 2D-G), a phenotype often seen in patients with large *GAP10* CNVs (DECIPHER). Interestingly, several individuals with large CNVs spanning *GAP10* exhibited proportionate short stature, short neck, small hands, and growth delay, phenotypes that may reflect conserved disruptions in axial elongation and morphogenesis (Table 1).

**Figure 2.**
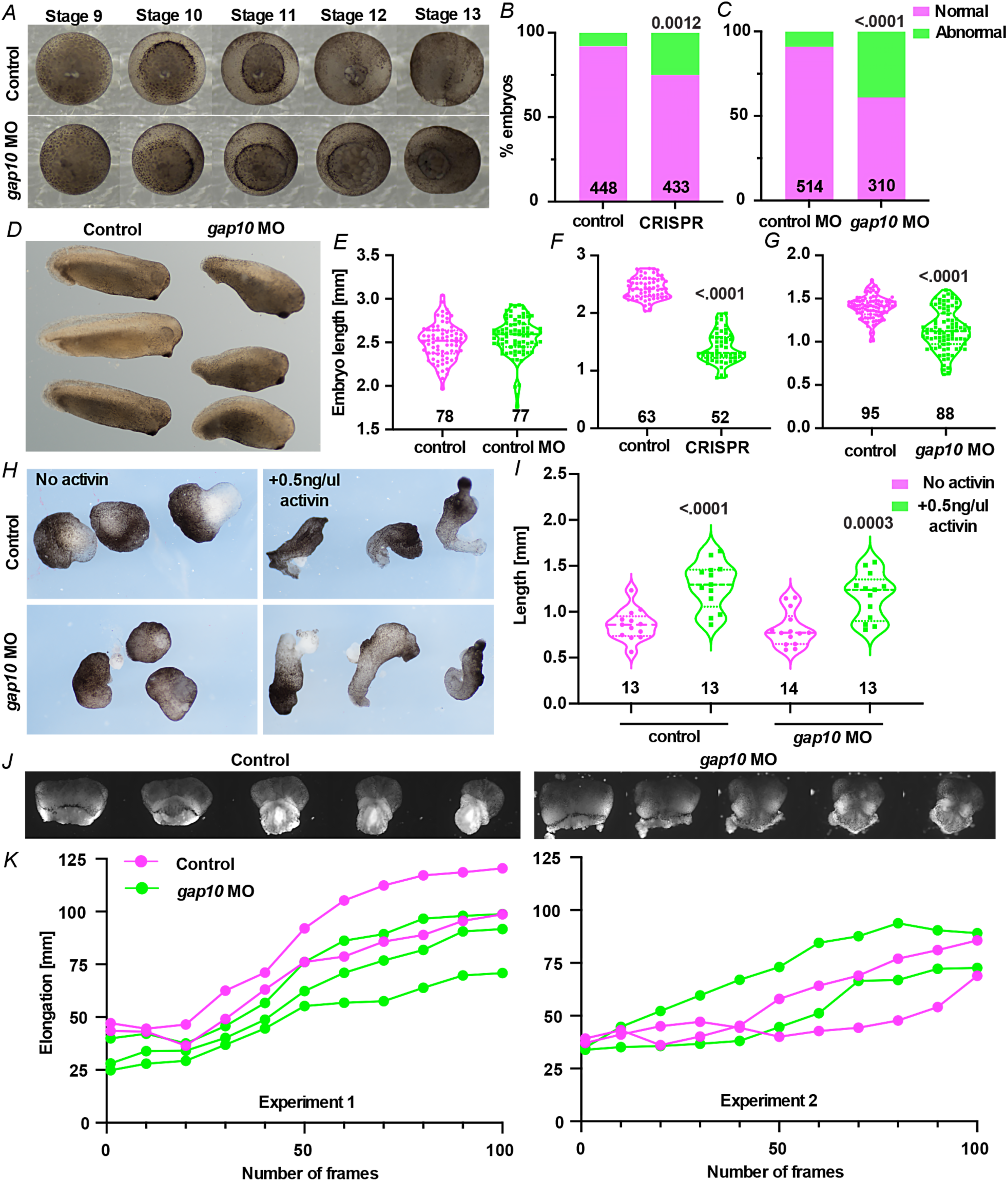
Gap10 is Required for Gastrulation and Body Axis Extension in *Xenopus* Embryos. **(A)** Montage of time lapse of control and *gap10* morphant embryos during gastrulation. **(B-C)** Quantification of delayed/incomplete blastopore closure in controls and *gap10*-deficient embryos, normal (magenta) versus abnormal (green). (n per group, p values as indicated). **(D)** Representative images of tailbud-stage embryos highlighting shortened body axis and dorsal curvature in *gap10* morphants. **(E-G)** Violin plots showing length and curvature measurements in control and *gap10*-deficient embryos at tailbud stages. (n per group, p values as indicated) **(H)** Activin-induced elongation assay showing explants with and without activin in control and *gap10*-deficient embryos. **(I)** Quantification of explant elongation with and without activin in control and *gap10*-deficient embryos (n per group, p values as indicated). **(J)** Time-lapse of morphogenetic movements in Keller’s sandwiches. **(K)** Quantification of the elongation of Keller’s sandwiches across two experiments.

To pinpoint the cause of axial shortening, we analyzed convergent extension (CE) movements in *gap10* morphant tissues^40^. Surprisingly, mesodermal CE appeared intact in activin-treated animal cap explants (which model mesodermal CE) from *gap10* morphants, elongated normally (Figure 2H, I)^41^. We next tested “Keller” explants, sandwiches of early gastrula dorsal marginal zone tissue used to observe CE *in vitro* (Figure 2J, K)^42,43^. *gap10* MO embryos appeared to have relatively normal extension, though not perfectly similar to controls. Thus, these results suggested that classical CE defects do not underlie the severe shortening of *gap10* morphants, pointing to an alternative mechanism for axial elongation failure. Notably, this phenotype parallels recent findings for the adherens junction protein ARVCF: loss of ARVCF disrupts whole-embryo body axis extension without affecting the extension of isolated explants, due to an inability to generate sufficient tissue-level force during morphogenesis^44^. By analogy, *gap10* may be required to integrate mechanical forces or cell-matrix interactions during embryonic elongation. In summary, *gap10* is essential for gastrulation movements and subsequent body axis formation.

### *Gap10* is Critical for Mucociliary Clearance and Multiciliogenesis

Beyond its role in LR development, motile cilia are also vital for airway mucociliary clearance. Indeed, patients with CHD, and especially heterotaxy, often suffer from chronic respiratory issues due to ciliary dysfunction^14–16^. Moreover, one patient with a GAP10 deletion was reported to experience respiratory distress (Table 1). While the etiology was not specified, this may reflect impaired mucociliary clearance. We therefore examined *gap10*’s function in multiciliated cells (MCCs) of the *Xenopus* embryonic skin, a well-established model for studying mucociliary clearance^45–47^. *gap10* morphants displayed severely impaired ciliary flow, as evidenced by particle tracking assays. In these assays, colored beads on the skin of morphant embryos showed little to no directed movement, whereas control embryos generated robust directional transport of beads through ciliary beating (Figure 3A-C)^47^. To determine whether this flow defect was caused by immotile cilia or abnormal ciliary morphology, we examined MCC differentiation using scanning electron microscopy (SEM) and immunofluorescence. MCCs in control embryos displayed a dense carpet of hundreds of elongated cilia, whereas *gap10* morphants showed a notable lack of motile cilia, and those that were present were often abnormally short (Figure 3D). Gap10 depletion did not affect the MCC number, suggesting it does not affect cell-fate specification (Figure 3E)^48^.

**Figure 3.**
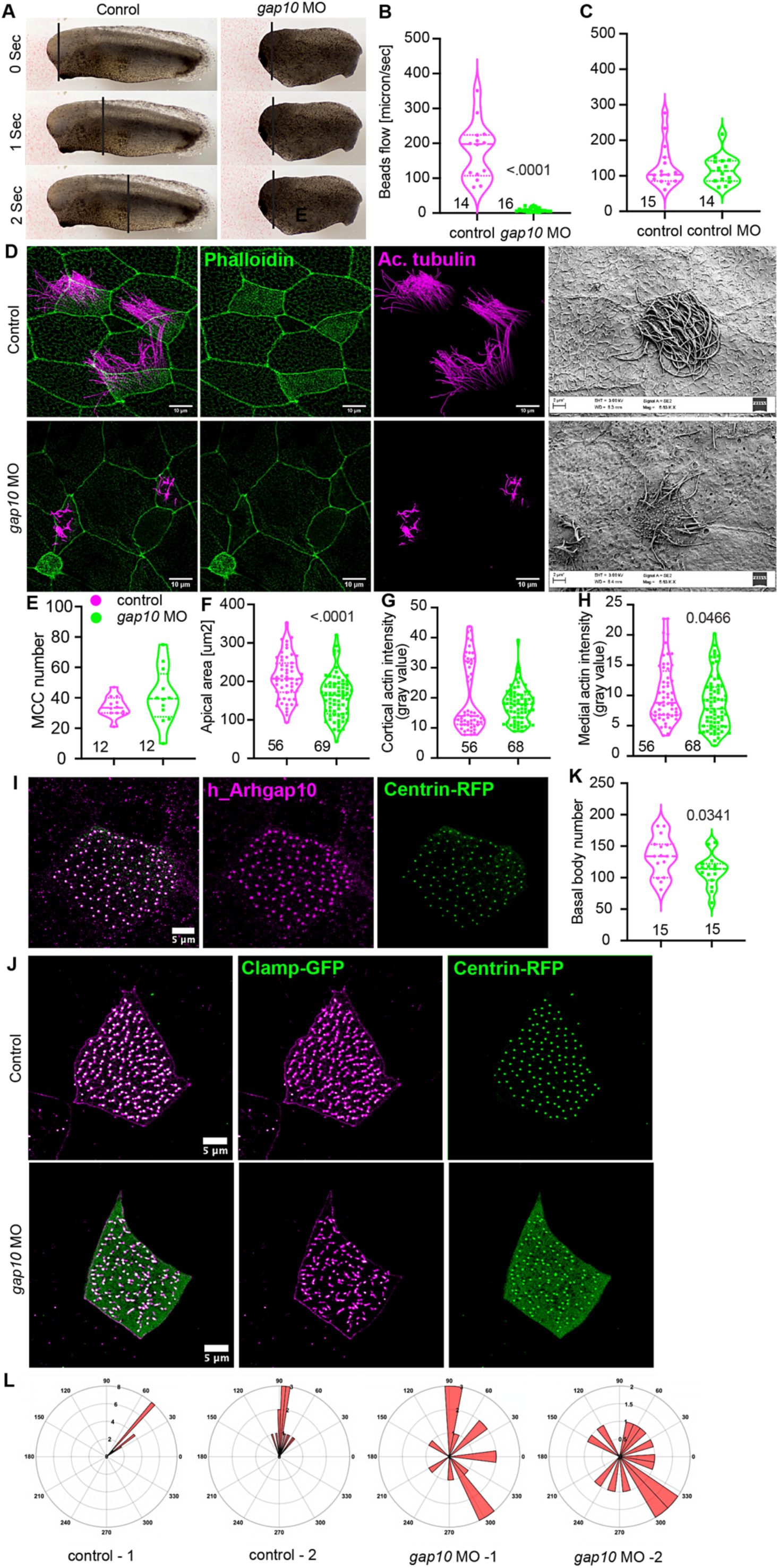
Gap10 Controls Mucociliary Differentiation, Actin Architecture, and Basal Body Organization in Multiciliated Cells. **(A)** Representative embryos showing lack of bead flow in *gap10* morphant compared to control. **(B-C)** Quantification of bead flow in uninjected controls and control MO and *gap10* MO knockdown embryos. (n per group, p values as indicated) **(D)** Phalloidin (green, F-actin) and acetylated tubulin (magenta, cilia) staining (maximum projections) of *Xenopus* embryonic skin showing multiciliated cells (MCCs) in control and *gap10* morphants (scale bars: 50 μm). Scanning electron micrographs illustrating MCC surface and ciliary defects in *gap10* morphants. **(E-H)** Quantification of MCC number, apical area, cortical and medial F-actin enrichment in MCCs of control and *gap10* morphants (n per group, p values as indicated). **(I)** Localization of hARHGAP10–GFP to basal bodies in MCCs, co-stained with centrin-RFP (basal bodies) (scale bars: 5 μm). **(J)** Loss of basal body spacing and polarity in MCCs of *gap10* MO knockdown embryos. **(K)** Analysis of basal body number in MCCs of control and *gap10* morphants (n per group, p values as indicated). **(L)** Analysis of basal body polarity by circular plots in two representative MCCs of control and *gap10* morphants.

During normal multiciliogenesis, MCCs undergo dramatic apical expansion and assemble an apical F-actin lattice that serves as a platform for basal body docking and spacing^47,49^. Phalloidin staining revealed that control MCCs possess an expansive meshwork of apical F-actin, whereas *gap10* morphants showed a fragmented or entirely absent apical actin lattice (Figure 3G-H). Consistently, the MCC apical surface in morphants failed to enlarge, remaining small and irregular (Figure 3F). In summary, Gap10 is essential for mucociliary clearance, as its absence causes defective MCC differentiation, characterized by an inability to expand the apical domain, failure to anchor basal bodies, and consequent loss of motile cilia needed to generate fluid flow.

### Human GAP10 Localizes to Basal Bodies and Is Essential for Basal Body Organization and Polarity

The loss of an apical actin network in Gap10-deficient MCCs closely phenocopies the effects of depleting WDR5, a chromatin modifier and a CHD/heterotaxy gene recently shown to have a non-transcriptional role at basal bodies of motile cilia^47^. WDR5 binds to basal bodies and acts as a scaffold to stabilize apical F-actin, and *Wdr5* knockdown likewise leads to loss of the actin lattice and failure of MCC apical expansion. Therefore, we investigated the localization of hGAP10 protein in MCCs. Exogenous GFP-tagged human GAP10 (hARHGAP10–GFP) expressed in *Xenopus* embryos localized prominently to basal bodies within MCCs (Figure 3I). This suggested that GAP10 might function at basal bodies. In normal MCCs, basal bodies are uniformly distributed across the enlarged apical surface and oriented in the same direction (planar polarity), which is crucial for coordinated ciliary beating (Figure 3J)^47^. In *gap10* morphants, basal bodies were fewer and no longer evenly spaced (Figure 3J, K). Instead, they clustered irregularly and failed to align in a uniform orientation, correlating with the reduction in cilia number and loss of directed flow (Figure 3J-L). Thus, *gap10* is required to establish proper basal body spacing and planar polarity in multiciliated cells.

### *Gap10* is Required for FAK Localization to Apical “Ciliary Adhesion” Complexes in MCCs

Recent research has identified specialized “ciliary adhesion” (CA) complexes at the base of motile cilia, consisting of classical focal adhesion proteins (FAK, Paxillin, Vinculin) repurposed to anchor basal bodies to the apical actin cytoskeleton^30^. GAP10 (also known as GRAF2) was initially identified as a GTPase-activating protein associated with FAK^20,21^. Notably, GRAF family proteins can bind the C-terminal domain of FAK through their SH3 domains and inhibit RhoA signaling by FAK to prevent excessive stress fiber formation^20,21,50^. We therefore hypothesized that *Gap10*’s role in cytoskeletal dynamics is through the focal adhesion signaling.

In control embryos expressing hGAP10–GFP, we observed that GAP10 puncta colocalized with FAK at the basal bodies (Figure 4A). To investigate whether Gap10 plays a role in recruiting FAK to these ciliary adhesion sites in multiciliated cells, we generated mosaic knockdowns (Figure 4B). Specifically, we injected FAK-RFP at one cell stage and *gap10* MO with membrane-GFP as a tracer in 1 out of 4 cells. Membrane-GFP also marks cilia, allowing us to distinguish between ciliated and non-ciliated cells. FAK-RFP is expected to be expressed in all MCCs. However, in *gap10* morphant MCCs (marked by membrane-GFP), FAK failed to localize properly; specifically, the FAK signal at basal bodies was greatly diminished or diffuse, indicating an inability to assemble the ciliary adhesion complexes when Gap10 is absent (Figure 4C, D). Note that basal bodies, although disorganized, are present in *gap10* morphants (Figure 3J), and thus, the reduced FAK signal at basal bodies in the morphants is not due to the absence of basal bodies. Thus, GAP10 recruits FAK to ciliary adhesion complexes, necessary for basal body anchoring and ciliogenesis. FAK normally functions as the mechanical linker between basal bodies and cortical actin, and FAK depletion is known to cause basal body migration/docking defects and ciliogenesis failure in MCCs^30^. This loss of FAK recruitment provides a direct explanation for the basal body disorganization and polarity phenotype described above. Gap10 appears to operate upstream of FAK, ensuring that FAK (and likely other focal adhesion proteins) are correctly positioned at the base of cilia to form the ciliary adhesion complexes that secure basal bodies to the actin cytoskeleton.

**Figure 4.**
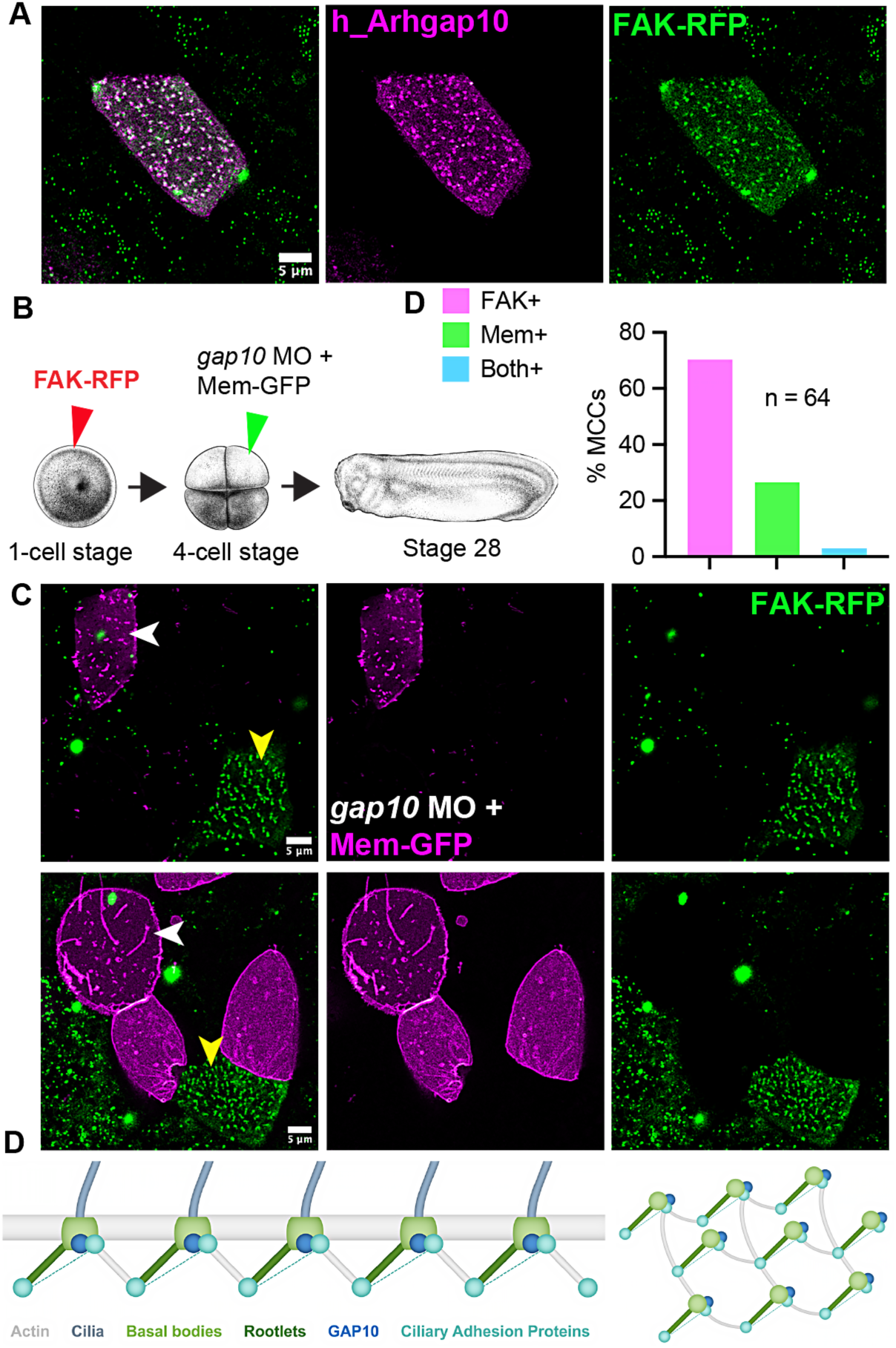
Gap10 is Required for FAK Localization at Ciliary Adhesion Complexes. **(A)** Confocal images of MCCs showing colocalization of hArhgap10 (magenta) and FAK-RFP (green) at basal bodies (scale bar: 5 μm). **(B)** Schematic of mosaic knockdown strategy: FAK-RFP injection at 1-cell stage; gap10 MO + membrane-GFP (tracer) at 4-cell stage for mosaic analysis. **(C)** Representative images showing loss of FAK-RFP signal at basal bodies (white/yellow arrowheads) in *gap10* morphant MCCs (scale bar: 5 μm).White arrowheads show membrane-GFP labeling defective cilia in *gap10* morphant MCCs and FAK-RFP is absent, whereas, MCCs lacking Membrane-GFP signal (yellow arrowheads) show FAK localizing to the bases of cilia. **(D)** Quantification of % MCCs with only FAK-RFP signal, or only Mem-GFP signal indicating loss of Gap10, or both signals. n is the number of MCCs. **(E)** Model illustrating the proposed role of GAP10 in recruiting FAK to ciliary adhesion complexes for basal body anchoring and actin network assembly.

GAP10, aka GRAF2, is a multi-domain Rho GTPase-activating protein belonging to the GRAF family. It contains an N-terminal BAR domain, a PH domain, and a C-terminal SH3 domain, which are architectural features that enable GRAF proteins to function as scaffolds, linking the membrane and the cytoskeleton^20,50^. Biochemically, GRAF2/GAP10 has GAP activity toward Rho family GTPases (notably RhoA and Cdc42) and can directly bind focal adhesion kinases via its SH3 domain^20^. These properties suggest a model in which GAP10 coordinates both the positioning of FAK and the local actin dynamics during multiciliogenesis. We propose that GAP10 is targeted to the apical cortex of MCCs, potentially through its BAR-PH domains sensing curved membrane or via interactions with actin-binding partners, where it helps assemble and stabilize the nascent ciliary adhesion complex^20,30,50^. There, GAP10 could tether FAK to the basal bodies, for example, by engaging proline-rich motifs on FAK (or its partners) via GAP10’s SH3 domain, thus anchoring FAK at the cilia base. In this way, GAP10 could orchestrate basal body attachment and apical cytoskeletal remodeling, which are essential for the development of multiciliated cells.

In conclusion, using an integrative *in vivo* functional genomics approach, we demonstrated that *gap10* is essential for cardiac development, embryonic elongation, and motile cilia function in *Xenopus*. Loss of Gap10 recapitulates patient phenotypes of CHD, short stature, and respiratory distress, mirroring the clinical consequences of *GAP10* CNVs. Our use of multiple, independent sgRNAs, along with similar morpholino results and clinical genetic evidence, strongly supports the specificity of the observed phenotypes. Collectively, these findings identify *ARHGAP10* as a novel CHD and ciliopathy gene, demonstrating how functional genomics can bridge rare human variants to disease mechanisms underlying developmental disorders.

## MATERIALS AND METHODS

### Animal husbandry and in vitro fertilization

Frogs (*Xenopus tropicalis*) were bred and housed in a vivarium using protocols (ACUC# 4295) approved by the University of Virginia Institutional Animal Care and Use Committee (IACUC). Embryos needed for experiments were generated using in vitro fertilization, as described previously^45,51^. Briefly, the testes from male frogs were crushed in 1× MBS (pH – 7.4) with 0.2% BSA and added to eggs obtained from the female frogs. After 3 min of incubation, freshly made 0.1× MBS (pH – 7.8) was added, and the eggs were incubated for 10 more minutes till contraction of the animal pole of the eggs was visible. The jelly coat was removed using 3% cysteine in 1/9× MR solution (pH 7.8–8.0) for 6 min. The embryos were microinjected with morpholino/RNA and raised at 25 °C or 28 °C until they reached appropriate stages for gastrulation, heart looping, fixation and staining, or bead flow analysis. Embryos were staged as described previously^52^. The fertilized embryos for microinjection were chosen randomly.

### CRISPR/Cas9 and Morpholino-Mediated Knockdown

To generate loss-of-function alleles, CRISPR/Cas9 editing was performed by microinjecting one-cell stage embryos with Cas9 protein (New England Biolabs) and sgRNAs (IDT, 400pg) targeting non-overlapping regions of *gap10*. sgRNA sequences were designed using CRISPRscan and are listed here, sg1 (Exon 1): GGCCAGGGGATTGGAATCAT, sg2 (Exon 1): GAGGATACGTGCCCATGAGG, and sg3 (Exon 4): GGGTTTCGGAAAGAGCAGTT. For knockdown experiments, translation-blocking antisense morpholino oligonucleotides (AGCCCCATGATTCCAATCCCCTGGC) were injected at the 1-cell stage (dose: 25 ng/embryo). Standard control morpholino and Cas9 only were used as negative controls for morpholino and CRISPR injections.

### Microinjection of mRNA and Tracer Constructs

Human ARHGAP10-GFP (NM_024605) (hGAP10-GFP, 400pg) and membrane-GFP (mGFP, 150pg) mRNAs were synthesized using the mMessage mMachine SP6 kit (Thermo Fisher) from linearized plasmids. mRNA was purified by column chromatography. mRNAs or morpholinos were injected at the one-cell or four-cell stage as indicated for mosaic or global perturbations. For mosaic knockdown, tracer mRNA (membrane-GFP) was coinjected for cell-autonomous analysis.

### In Situ Hybridization

Whole-mount in situ hybridization was performed using digoxigenin-labeled antisense probes for *Pitx2* and *Coco/Dand5,* following standard protocols^32^. Embryos were fixed in MEMFA (100 mM MOPS, 2 mM EGTA, 1 mM MgSO₄, 3.7% formaldehyde) at relevant stages. Probe hybridization and detection used alkaline phosphatase-conjugated anti-DIG antibody (Roche) with BM Purple substrate. LR marker scoring was performed blinded to injection status.

### Immunofluorescence and Imaging

Embryos were fixed in 4% paraformaldehyde at appropriate stages. After PBST washes (1× PBS + 0.2% Triton X-100) and blocking (3% BSA in PBST), samples were incubated with primary antibodies: mouse anti-acetylated tubulin (1:1000; Sigma), rabbit anti-GFP (1:500; Abcam), and phalloidin-AlexaFluor conjugates (1:400; Thermo Fisher) for F-actin. Secondary antibodies were AlexaFluor-conjugated (1:1000; Invitrogen). Embryos were mounted in ProLong Gold (Invitrogen) and imaged on a Leica SP8 confocal microscope (40× oil, 1.3 NA). Images were processed and quantified in Fiji/ImageJ and assembled in Adobe Illustrator software.

### Scanning Electron Microscopy

For MCC surface imaging, embryos were fixed in 2.5% glutaraldehyde in 0.1 M cacodylate buffer, post-fixed in 1% osmium tetroxide, dehydrated, and critical-point dried. Samples were sputter-coated with gold-palladium and imaged on a Zeiss Sigma FE-SEM.

### Ciliary Flow and Bead Tracking Assay

Embryos were raised to stage 28 and anesthetized with benzocaine (0.05% in 1/9x MR). One microliter of 5.19 µM red beads (Bangs Laboratories, DSCR006) was placed at the anterior end of the embryo and visualized under a dissecting scope. Bead movement was imaged by time-lapse under a stereomicroscope.

### Basal Body and Planar Polarity Analysis

Basal bodies were labeled by expression of centrin-RFP (100pg) or Clamp-GFP (150pg) mRNAs. Planar polarity was quantified by measuring basal body orientation angles in MCCs using ImageJ plugins. Apical area and spacing were measured on maximum-intensity projections of confocal stacks.

### Patient CNV and Clinical Data

Patient data were obtained from the DECIPHER and previously published manuscripts. Coordinates, gene content, and clinical phenotypes were curated (see Table 1).

### Statistical Analysis

Statistical significance was assessed in GraphPad Prism using Fisher’s exact test for categorical data (e.g., heart looping), and unpaired t-tests, or Mann–Whitney U tests for continuous variables as appropriate. Sample sizes (n) and number of biological replicates are indicated in figure legends.

### Data and Code Availability

All data supporting this study are provided within the article and Supplementary Material. Plasmids and reagents are available from the corresponding author upon reasonable request.

## ACKNOWLEDGMENTS

We thank Dr. Karen Hirschi for providing access to the confocal microscope. We are grateful for the NIH grants: NIGMS R35GM146856, NIH R03HD112688, and a grant from the Saving Tiny Hearts Society, awarded to Saurabh Kulkarni, as well as R35GM131865, awarded to Dr. DeSimone.

## AUTHOR CONTRIBUTIONS

ES: Investigation, Visualization, and data analysis.

SH: Figure preparations, manuscript writing, and revisions.

DS: Keller sandwich experiments, manuscript feedback, and revisions.

DD: Manuscript feedback and revisions.

SSK: Conceptualization, Methodology development, Investigation, Visualization, Supervision, and Manuscript writing and revisions.

## Inheritance / Genotype

De novo (unconfirmed parentage)

Heterozygous

De novo (unconfirmed parentage)

Heterozygous

Inherited from normal parent

Heterozygous

De novo (unconfirmed parentage)

Heterozygous

De novo, mosaic

Heterozygous

De novo (unconfirmed parentage)

Heterozygous

De novo (unconfirmed parentage)

Heterozygous

Unknown

Heterozygous

## Phenotypes

Ventricular septal defect

Atrial septal defect; Atrioventricular canal defect

Proportionate short stature

Growth delay

Short foot; Short neck; Small hand

Abnormality of prenatal development or birth; Atrial septal defect; Respiratory distress

Atrial septal defect; Proportionate short stature; Ventricular septal defect

Abnormal heart morphology

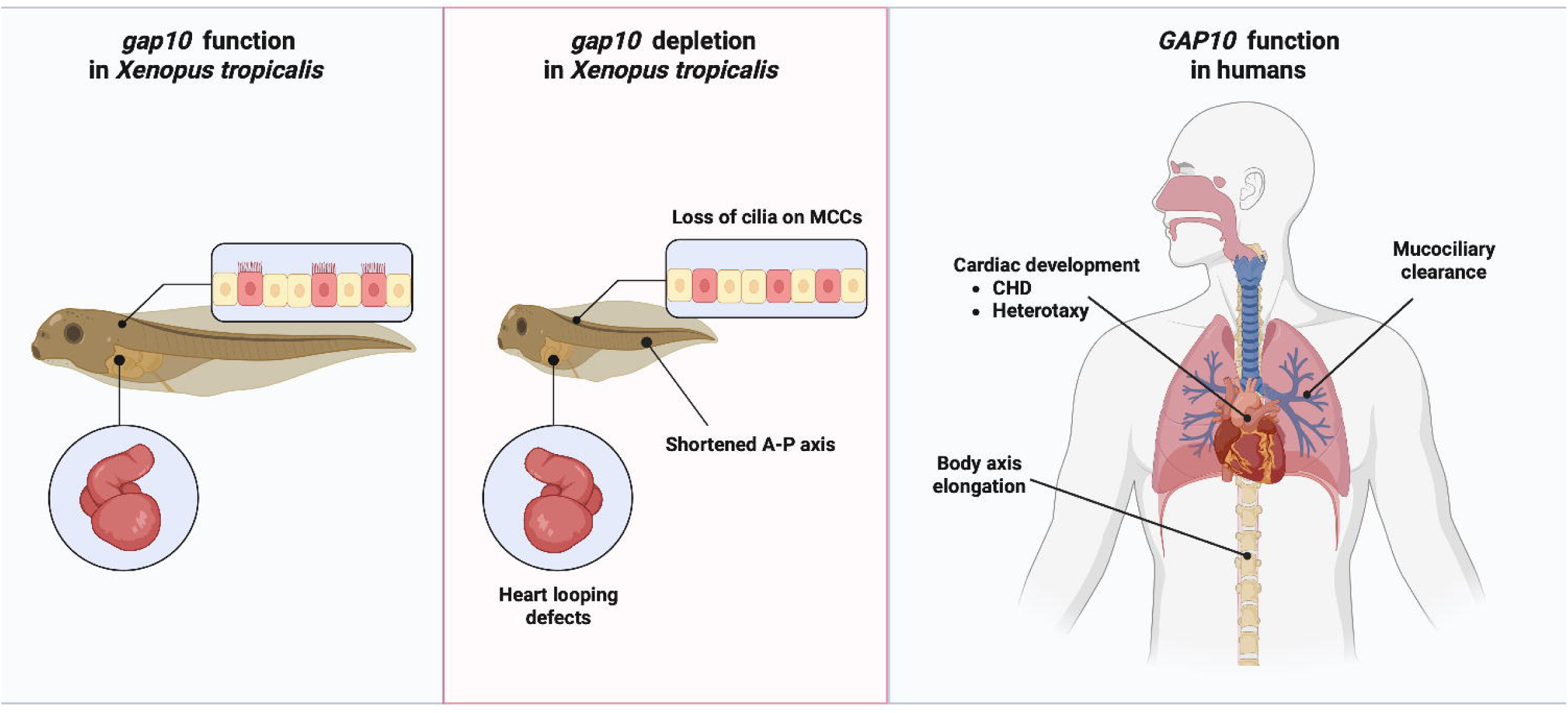

## Notes

### Competing Interest Statement

The authors have declared no competing interest.

